# High-affinity biomolecular interactions are modulated by low-affinity binders

**DOI:** 10.1101/2023.03.31.535017

**Authors:** S. Mukundan, Girish Deshpande, M.S. Madhusudhan

## Abstract

Molecular interactions play a central role in all biological processes. The strength of these interactions is often characterized by their dissociation constants (*K_D_*). The high affinity interactions (*K_D_* ≤ 10^-8^ M) are crucial for the proper execution of cellular processes and are thus extensively investigated. Detailed molecular and biochemical analyses of such high affinity interactions have lent considerable support to the concept of binary on/off switches in different biological contexts. However, such studies have typically discounted the presence of low-affinity binders (*K_D_* > 10^-5^ M) in the cellular environment. In this study, we have assessed the potential influence of such low affinity binders on high affinity interactions. By employing Gillespie stochastic simulations as well as continuous methods, we demonstrate that the presence of low-affinity binders can indeed alter the kinetics and the steady state of high-affinity interactions. We refer to this effect as ‘herd regulation’ and have evaluated its possible impact in two different contexts including sex determination in *Drosophila melanogaster* and in signalling systems that employ molecular thresholds. Lastly, we suggest an ‘in vitro’ experimental strategy to validate herd regulation. Based on these analyses, we propose that low-affinity binders are likely prevalent in different biological contexts where the outcomes are contingent upon threshold value determinants and thus potentially impact their homeostatic regulation.

## 1 Introduction

Key to all biological processes are biomolecular interactions. Networks of these interactions are organised into pathways, that in many instances, intersect (Huh et al., 2003; Kanehisa et al., 2022). These pathways usually consist of tight binding interactions. In a cellular context, these interactions do not occur in isolation but in the presence of a plethora of other molecules and pathways. This, in turn, ensures that biomolecules can promiscuously interact with multiple partners with varying degrees of affinity. It stands to reason that some of these interactions and their interaction strengths have evolved to be integral parts of signalling pathways.

In this study, we have analysed the influence of specific but low affinity binders on tight binding high affinity interactions. The strengths of these interactions are reported in terms of their dissociation constants, *K_D_*. *K_D_* is the equilibrium constant of the reverse reaction given by the ratio of the reverse and forward rate constants. Typically, members of a pathway would interact with their partners, with *K_D_* values in the range of 10^-6^ M to 10^-14^ M (Hu, 2017; Perkins et al., 2010), with lower values corresponding to stronger binding. However, the cellular environment consists of a variety of interactors with variable affinities. In addition to its specific high affinity binding partner(s), a biomolecule could encounter a cohort of other binders with *K_D_* values > 10^-6^ M present in the vicinity. Many of these lower affinity interactions could be energetically favourable (negative ΔG) and thus are competent to sequester the reactants from the high affinity reaction.

Studies of biological processes usually focus almost exclusively on the important tight binding interactions. This is in part because these canonical interactions are considered to be central to cellular processes. Moreover, experimental techniques mostly detect strong interactors (Galaway and Wright, 2020; Jiang and Barclay, 2009; Li et al., 2014). Here, we have examined how these high affinity (tight binding) interactions could be altered and/or modulated in the presence of several specific low affinity binders. For this study, low affinity interactors were conservatively considered to have *K_D_* values of 2 to 5 orders of magnitude less favourable than their stronger counterparts. We also vary the concentration of the low affinity binders to be in the range of 1X to 10X to that of the tight binding partners. This is a conservative estimate as the cellular milieu may contain a higher concentration of the low affinity binding partners, mechanisms for local enrichment of binding partners notwithstanding.

Using these values of binding strengths and relative concentrations, we investigated the combined effect of such weak binding partners on the kinetics and equilibrium of a tight binding interaction. We computed these effects using Gillespie stochastic simulations (Gillespie, 1977, 2013) and also replicated them with a system of differential equations. For instance, consider the following interaction.

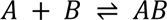

Here, the reactants *A* and *B* interact tightly to form the product *AB*. Let *A_i_* and *B_i_* represent the cohort of specific low affinity binders of *B* and *A* respectively. We have studied how the kinetics and steady-state changes with varying levels and binding affinities of *A_i_* and *B_i_* in the cases where – i) there are low affinity binders for either *A* or *B*, ii) both reactants have low affinity binders, and iii) there are low affinity binders to *A* and *B* and the *A_i_*’s and *B_i_*’s can interact with each other. We have coined the term herd regulation to represent the model of low-affinity binders interfering with high affinity interactions. Intuitively, this would be possible when the low-affinity interactions are in sizeable quantities (herds).

In a cellular context, such interactions should be prevalent as the low affinity binders are likely widespread. Their presence could modulate molecular interactions and hence biological activities. Here, we have modelled the process of sex determination in *Drosophila melanogaster* and signalling thresholds of biomolecular processes, two processes whose outcomes are dependent on thresholds.

*Sex-lethal* (*Sxl*), a master-switch gene, which determines the female identity, is turned on by the doubling of the X chromosome signal in females (Cline and Meyer, 1996). Analysis of mutations in X-chromosome-linked activators has confirmed that activation of early embryonic promoter of *Sxl* (*Sxl-Pe*) is critical in females and thus, underlies the binary fate choice. By contrast, strong repressors capable of affecting sex determination i.e. sex-specific identity and/or viability, have not been identified (Mahadeveraju et al., 2020; Salz and Erickson, 2010). Loss of function mutations in such repressors would result in anomalous activation of *Sxl-Pe* in males, which in turn, can induce male-to-female transformation or lethality. Here, we hypothesise that such repressors are a set of low affinity binders. A single mutation, typically isolated during genetic screens, however, would not elicit strong phenotypic consequences (e.g. sex-specific lethality) used as a conventional and stringent criteria for selection. To examine the possible impact of such repressors, we have modelled this system considering *A* as the switch gene, *B* as the X chromosome signal and *A_i_* and *B_i_* as repressors. We demonstrate that our simulation strategy yields outputs that are consistent with previously reported experimental findings.

Successful activation of biological signalling pathways often requires signal strength to attain or surpass a defined threshold value. Such systems involve the conversion of a linear change in the signal, to a binary on/off state. Morphogen gradients leading to cellular patterning in a variety of developmental contexts constitute one such example (Wartlick et al., 2009). Although such cases are prevalent in biological systems, precise molecular mechanisms underlying these decisions are not always fully understood. Previous modelling attempts at elucidating the mechanisms have employed models that depend on cellular compartments and/or promiscuous binders (Iyer et al., 2022). In this study, we provide a simple yet broadly applicable mechanism that explains such thresholds using low affinity binders.

## 2 Results

### 1.1 The effect of low affinity competitors on strong interactions

#### 1.1.1 Only one of the strong interactors has competition

To start with, we examined the effect of low affinity interactors on one of the components of a tight binding complex, i.e., the effect of *B_i_* on the formation of the complex *AB* (see equations E1 and E2). The outcome of these reactions is intuitive but serves the crucial purpose of creating a control case for subsequent sections. *AB* (dissociation constant *K_D_* = 10^-8^ M) is 3 orders of magnitude stronger than *AB_i_* (dissociation constant *K_D_i_* = 10^-5^ M). We have monitored the effect of two parameters, initial levels of the low affinity interactors, *B_i0_*, and their binding affinities, *K_D_i_*. We analysed the kinetics and steady states of these reactions using the Gillespie stochastic simulation algorithm (Gillespie, 2013).

First, we studied the effect of *B_i0_* levels. The simulation was set up with 500 copies of *A* and *B* and with *K_D_* = 10^-8^ M. Kinetics of reactions with different levels of *B_i0_*, ranging from 0 to 10000 were analyzed at *K_D_i_* = 10^-5^ M. At the beginning of the reaction the entities *A*, *B* and *B_i_* are present as free monomers. Almost instantaneously, all the monomers of *A* are bound to either *B* or *B_i_*, proportional to their relative abundance. As expected, the larger the value of *B_i0_*, the longer the system takes to equilibrate (figure 1A). This is because the forward rate constants of the formation of *AB* and *AB_i_* are assumed to be equivalent. Consequently, the levels of *AB_i_* may be transiently higher than that of *AB* (see figure A6 panel A for the case where the reverse rate constant is equal). *AB_i_* levels decrease while *AB* levels increase since *AB_i_* are more likely to dissociate than *AB.* (Figure 1A). As a result, the average time taken (from 50 replicates) for the formation of *AB* (90% of the maximum possible level, rise time) increases with rising levels of *B_i0_* (figure 1B).

**Figure 1:**
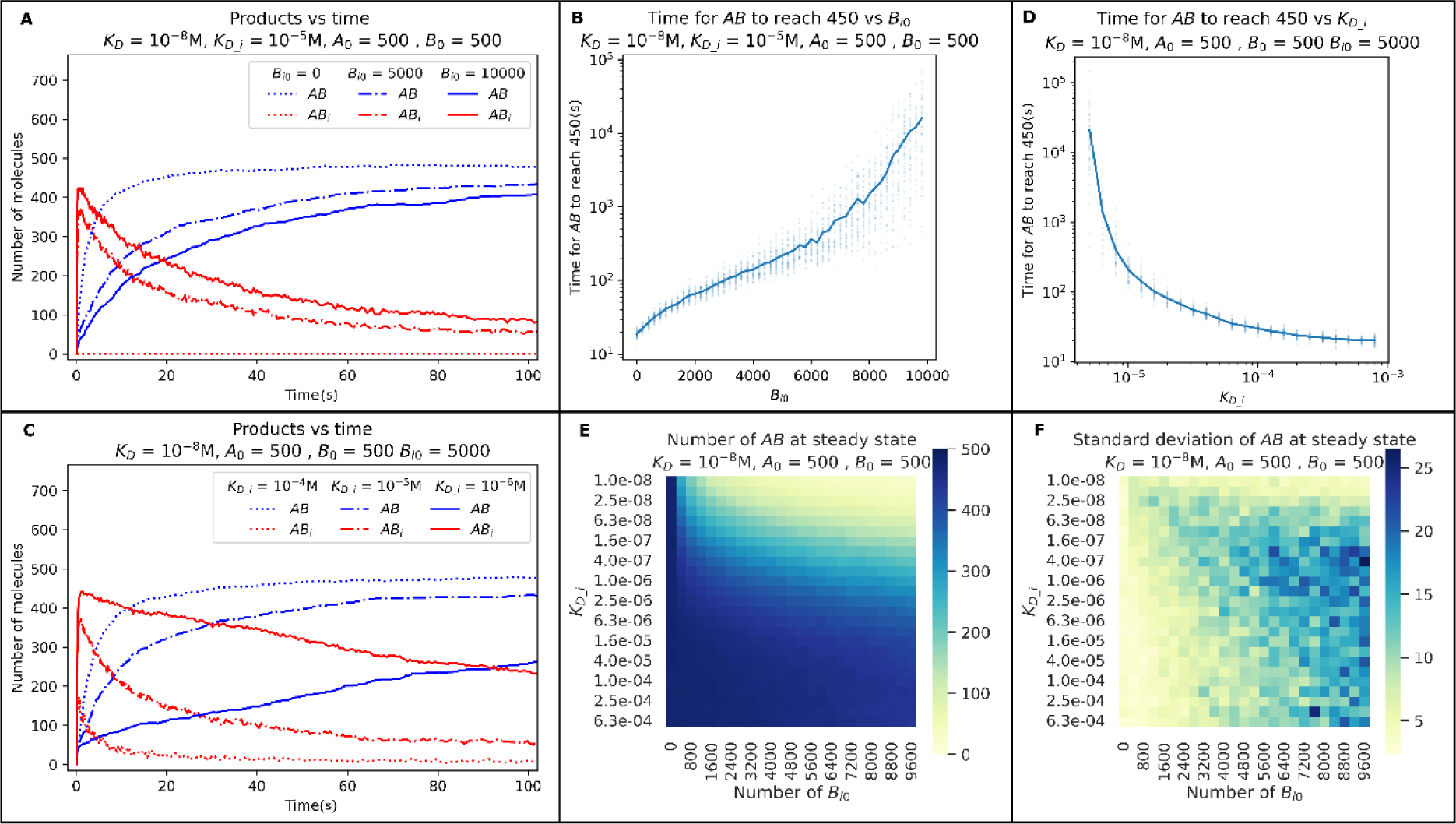
The dependence of complex formation on the starting levels and dissociation constants of the low affinity binder *B_i_*. The kinetics of *AB* (blue lines) and *AB_i_* (red lines) formation at different *B_i0_* and *K_D_i_* (panels **A** and **C** respectively). The effect of *B_i0_* and *K_D_i_* on the number of *AB* at steady state (average of 25 replicates per box in panel **E**) with the standard deviations in panel **C**. The time taken for *AB* to reach 450 molecules (90% of the maximum possible value of 500) at different *B_i0_* and *K_D_i_* levels (panels **B** and **D** respectively). Panels **A** and **C** show single trajectories taken from triplicate simulations. The lines in panels **B** and **D** are averages over 50 replicates with individual data points shown as scatter plots. The parameters used are shown above the figures.

We also examined the effect of variation in the dissociation constants of low-affinity interactors, on the formation of *AB.* This was done by varying *K_D_i_* from 10^-4^ M to 10^-6^ M while maintaining a *B_i0_* level of 5000. Again, as expected, decreasing *K_D_i_* (increasing the binding strength of the low affinity binder) decreases the rise time of *AB* (figure 1C). When *K_D_i_* is greater than 10^-4^ M, the formation of AB complex breaches the 90% level almost instantaneously. At the other extreme, when *K_D_i_* is smaller than 10^-6^ M (getting closer to K_D_), intuitively, the rise time of the AB complex tends to infinity (figure 1D).

To calculate the amount of *AB* found at equilibrium, we have analysed the steady state levels of the AB complex at 625 individual combinations of 25 *B_i0_* levels ranging from 0 to 9600 and 25 *K_D_i_* values ranging from 10^-3^ M to 10^-8^ M, each replicated 25 times. We observed that AB is the major product as long as *K_D_i_* is greater than 10^-7^ (figure 1E). However, *B_i0_* levels start to affect the amount of *AB* formed beyond 5000 molecules. Interestingly, variance in the outcome of the reaction also increases with increasing *B_i0_* and increasing *K_D_i_* (figure 1F). This is quite counterintuitive as one would expect the variance from Gillespie stochastic simulations to decrease with an increase in the size of the system. The two regions in figure 1F with low variance are either dominated by *AB* or *AB_i_* and the ‘twilight zone’ where both complexes are found in comparable amounts has high variations.

Since low affinity interactors acting on one component of the high affinity interaction can significantly impact the rate of high affinity interaction, we sought to assess the effect of low affinity interactors on both the high affinity binders.

#### 1.1.1 Both strong interactors face competition

In addition to low affinity interactors of *A* (*B_i_*), we now also consider the presence of interactors of *B* (*A_i_*) in the reaction. *A_i_* interacts with *B* and forms the complex *A_i_B* (see equations E1, E2 and E3). The dissociation constants of *AB_i_* and *A_i_B* are both described by a single term, *K_D_i_*, as we consider them to be equivalent. For this reason, all the results only describe/show *AB_i_*, while the equivalent *A_i_B* is not shown. As in the previous section, the binding affinity of the tight binder (*K_D_*) is 3 orders of magnitude stronger than *K_D_i_*. We analyzed the effect of varying the starting levels of *B_i_* and *A_i_*, as well as their binding affinities *K_D_i_* on *AB* formation using Gillespie stochastic simulations.

We first examined the effect of starting levels of *B_i_* and *A_i_* on *AB* formation. Simulations were set up with 500 copies of *A* and *B* and with *K_D_* = 10^-8^ M. The kinetics of reactions with different levels of *A_i0_* and *B_i0_*, ranging from 0 to 10000 were analyzed at *K_D_i_* = 10^-5^ M. We observed that the presence of low affinity binders increases the rise time of *AB* (figure 2A). The time taken for *AB* to equilibrate is an order of magnitude higher than what we noticed when only *B_i_* was present in the system (figures 1A and 2A). Also, the rise time increases exponentially with increasing values of *A_i0_* and *B_i0_* (figure 2B). These results suggest a non-linear effect arising from the addition of one more low affinity binder.

**Figure 2:**
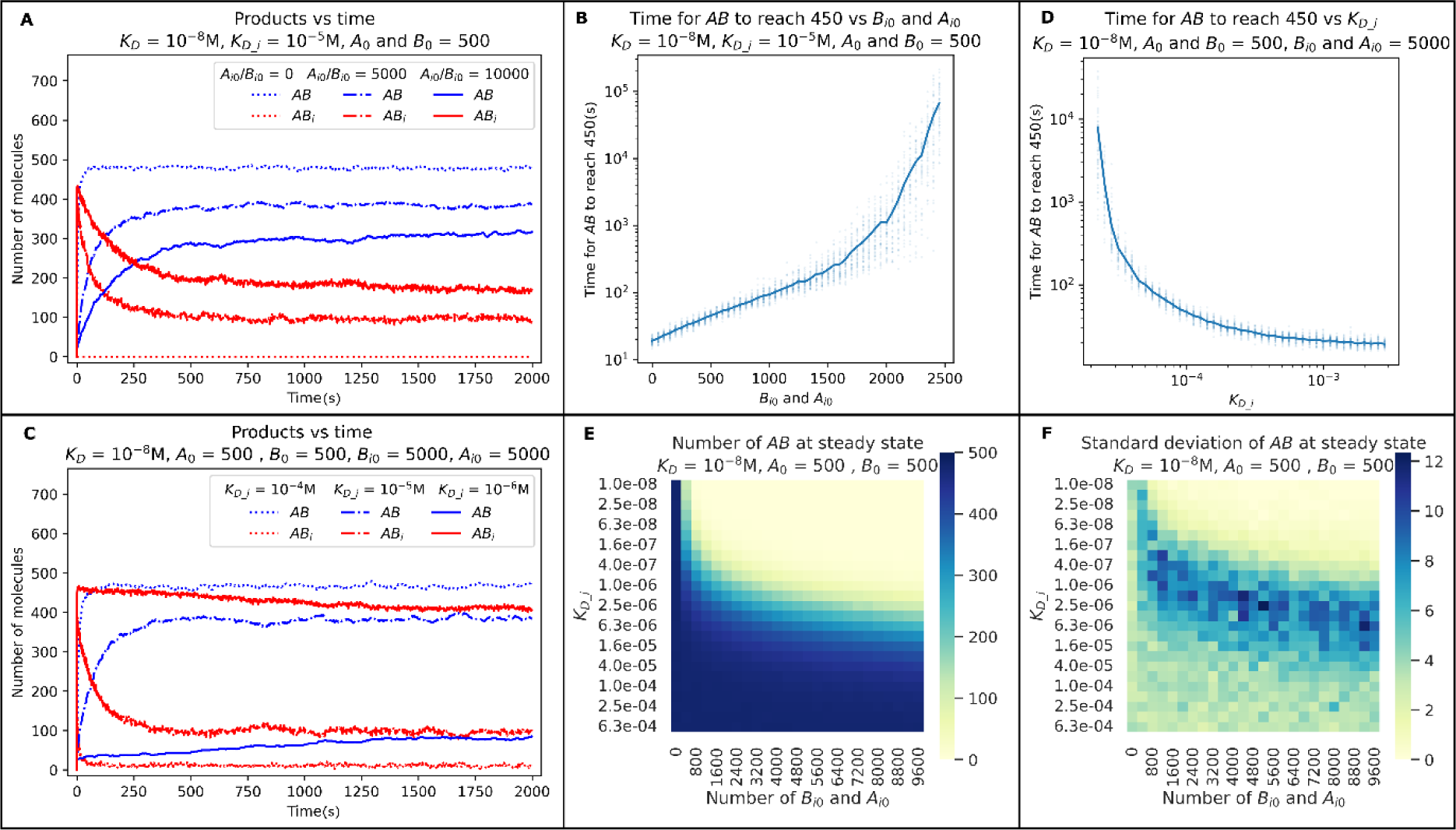
The dependence of complex formation on the starting number and dissociation constant of low affinity binders *B_i_* and *A_i_*. The kinetics of *AB* (blue lines) and *AB_i_* (red lines) formation at different starting levels and dissociation constants of *B_i_* and *A_i_* (panels **A** and **C** respectively). The effect of different starting levels and dissociation constants of *B_i_* and *A_i_* on the *AB* levels at steady state (average of 25 replicates per box in panel **E** and the standard deviations in panel **F**). The time taken for *AB* to reach 450 (90% of the maximum possible value of 500, black dotted line in panels **A** and **C**) at different starting levels and dissociation constants of *B_i_* and *A_i_* (panels **B** and **D** respectively). Panels **A** and **C** represent single trajectories taken from triplicate simulations. The lines in panels **B** and **D** are averages over 50 replicates. The parameters used are shown above the figures.

We then investigated the effect of *K_D_i_* on *AB* formation at values ranging from 10^-4^ M to 10^-6^ M. Decreasing *K_D_i_* increases the time taken for *AB* to equilibrate (figure 2C). Surprisingly, at *K_D_i_* = 10^-6^ M (two orders of magnitude weaker than AB), the reaction seems to prefer the formation of the low affinity complex (figure 2C). We also found that the rise time exponentially increases with decreasing dissociation constant (figure 2D).

To investigate this further, we analysed the steady state of these reactions using 625 different combinations of 25 *A_i0_* and *B_i0_* levels ranging from 0 to 9600 and 25 *K_D_i_* values ranging from 10^-3^ M to 10^-8^ M. AB is not the major product if the number of *A_i0_* and *B_i0_* are both above 3000 and *K_D_i_* is below 10^-6^ M (figure 2E). This is despite a *K_D_* of 10^-8^ M and with 500 copies of both *A* and *B*. The variability in outcome for reactions within this range is also low. Remarkably, the variability in *AB* levels at steady state is comparatively higher betwixt the areas that completely prefer *AB* or *AB_i_* (figure 2F). Thus, the simultaneous presence of the weak binders, *A_i0_* and *B_i0_,* in the system, has a greater impact on the formation of *AB*, than when only one weak binder is present. This enhancement is sufficient to allow the weaker complex to become the major product. Note that adding *A_i_* to the system does not increase the concentration of one binder as *B_i_* and *A_i_* binds specifically to *A* and *B,* albeit with low affinity. Henceforth, we refer to this effect of weak binders as *herd regulation*.

The presence of monomeric *A_i_* and *B_i_* in the system is likely responsible for the sharp decline in *AB* formation. This is because monomeric *A* and *B* formed by the dissociation of *AB_i_* or *A_i_B* have a higher chance of interacting with *B_i_* and *A_i_* respectively than with another tight binding monomer. To assess if this is true, we have analyzed the outcome when free *A_i_* and *B_i_* can interact with one another to produce *A_i_B_i_*, a system with minimal amount of monomeric *A_i_* and *B_i_*.

#### 1.1.2 Interaction amongst the low affinity competitors decreases herd regulation

We investigated the effect of low affinity interactors *A_i_* and *B_i_* when they can bind to each other to give rise to the reaction product *A_i_B_i_* (E4). The dissociation constant of *A_i_B_i_* is denoted by *K_D_cross_*. We set the dissociation constant for *AB_i_* and *A_i_B* as 10^-6^ M, a value at which herd regulation favours the low affinity products (figure 2C). We simulated the effect of cross reaction between *A_i_* and *B_i_* on the kinetics of this system using Gillespie stochastic simulations.

We found that decreasing *K_D_cross_* (increasing cross reaction strength) increases the rate of *AB* formation (figure 3A). This is intuitive as it decreases the net concentration of low affinity monomers present. But it is noteworthy that in this section we are expanding on a particular case from the previous section where the low affinity binders were dominant even with 2 orders of magnitude weaker binding affinity. Also, note that even with a strong cross-reaction (compared to *K_D_i_*) the system prefers weak binding complexes. Also, the time taken for *AB* to reach 90% saturation decreases with decreasing *K_D_cross_* (figure 3B). Thus, stronger *A_i_B_i_* interactions favour more active product formation by sequestering the monomeric *A_i_* and *B_i_* as *A_i_B_i_*, allowing the lower probability dissociation reactions to occur and leading to the formation of the tight binding complex.

**Figure 3:**
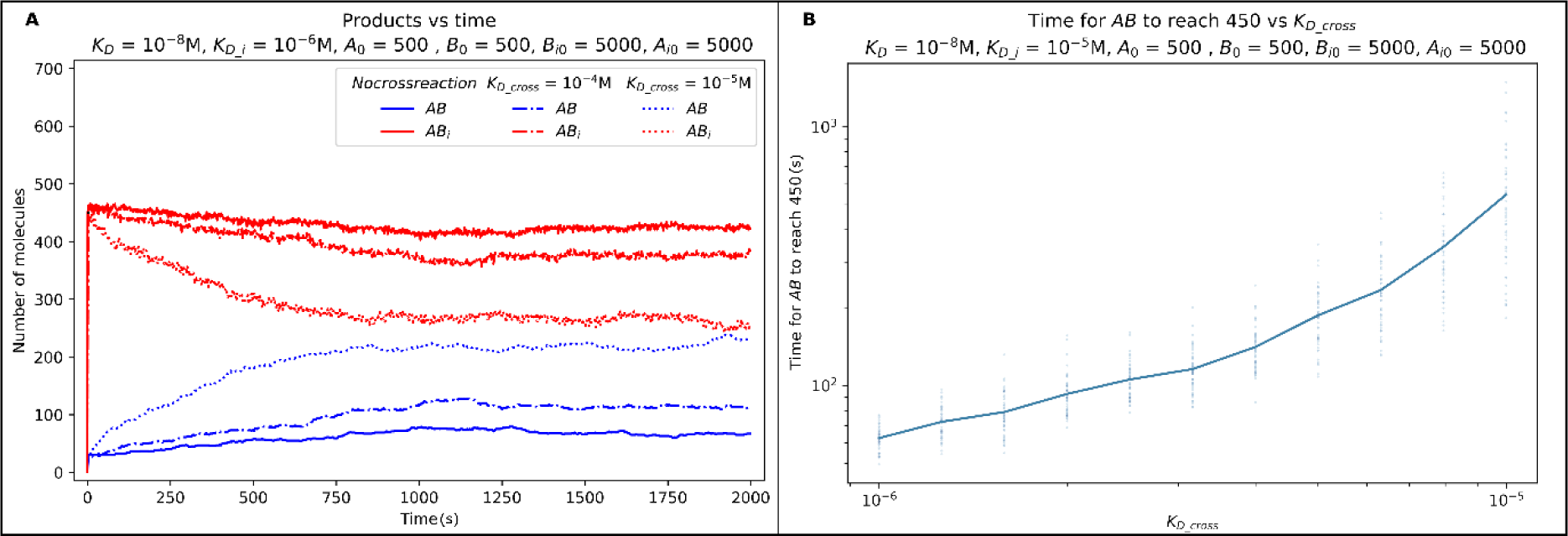
The effect of cross-reaction between low affinity inhibitors on herd regulation using Gillespie stochastic simulations. The kinetics of *AB* (blue) and *AB_i_* (Red) complexes at cross-reaction dissociation constants (*K_D_cross_*) 10^-4^M and 10^-5^M (panel **A**). Panel **B** shows the effect of *K_D_cross_* on the time taken for *AB* to reach 450 (90% of the maximum possible value of 500, black dotted line in panel **A**. One of 3 replicates are plotted in panel A. The plot in Panel **B** shows averages of 50 replicates (shown as scatter). Starting values of *A*, *B* = 500, *K_D_* = 10^-8^M and *K_D_i_* = 10^-6^M. Parameters shown above the figures.

Since we established that low affinity binders can affect the kinetics and equilibrium of tight binding interactions we now aim to formulate a general model that describes this effect (herd regulation).

#### 1.1.3 General model for describing herd regulation with multiple interactors

In this section, we have formulated a general model that describes the kinetics and equilibrium of a herd regulated system based on sets of ordinary differential equations.

Consider the *N* bimolecular association reactions simultaneously happening in the same reaction volume.

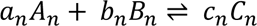

Here *a*, *b* and *c* are the stoichiometric coefficients of *A*, *B* and *C* respectively. And *n* denotes the individual reaction involved in the process. For every *n* ∈ 1,2…*N*

The net rate of formation of complexes for these reactions is defined as the sum of forward and reverse rates at any given time point. The following set of equations defined the rates of formation of each complex.

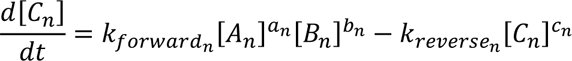

Here [*A*_*n*_], [*B*_*n*_] and [*C*_*n*_] represent the concentration of the respective entities at a given point in time and, *k*_*forward_n_*_ and *k*_*reverse_n_*_ represent the forward and reverse (on and off) rate constants for a reaction.

The rate of change of the complex is 0 at equilibrium and *K*_*D*_ is the ratio of the reverse and forward rate constants. Thus, the above equation simplifies to the following set of equations that describe the system at equilibrium.

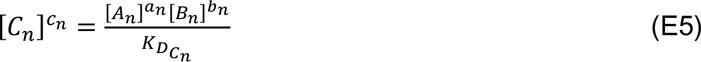

The equilibrium states can be determined by solving the above set of equations.

The states of *A*_*n*_ and *B*_*n*_ at any given time can be defined in terms of their initial concentrations, the concentration of complexes that they are part of at a given time and their stoichiometry.

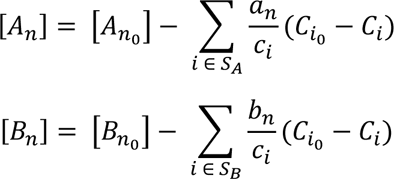

Here *S_A_* is the set of complexes that consume *A_n_* and *S*_*B*_ is the set of complexes that consume *B_n_*. The concentration terms with subscript 0 are the initial concentrations of the respective entities.

This model can be used to generate equations that describe specific cases of herd regulation. For example, the following set of equations describes the system of reactions used in the study i.e. E1, E2, E3 and E4 (methods section 4.1).

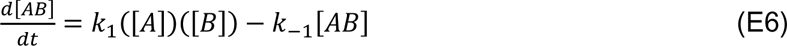

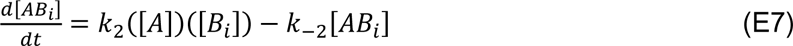

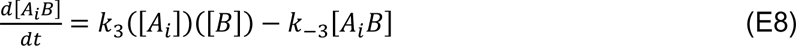

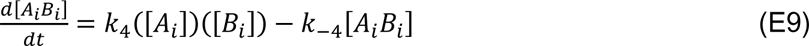

The equilibrium states can be deduced by solving,

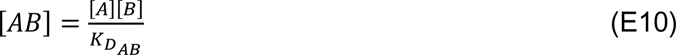

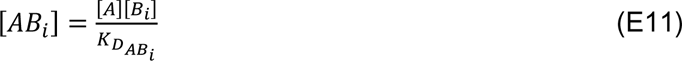

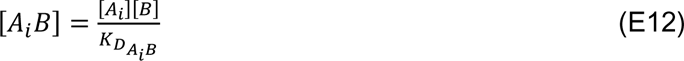

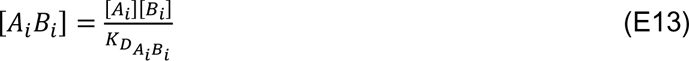

Since the reactants are shared by the reactions *A*, *B*, *A_i_* and *B_i_* are affected by multiple reactions and can be expressed as shown below.

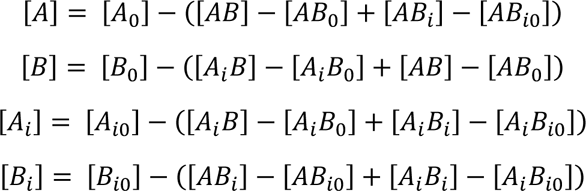

Here the concentrations with a subscript of 0 imply the initial concentration

Using equations E6-E13 we computed the trajectories and equilibrium states for herd regulation with i) only *B_i_* in section 8.4.1, ii) with *B_i_* and A*_i_* in section 8.4.2 and iii) where *B_i_* and *A_i_* cross-react in section 8.4.3. The results from this continuous model closely match the results obtained from Gillespie stochastic simulations.

Note that the equilibrium state only depends on the dissociation constant and is completely independent of the forward and reverse rate constants. Thus, our assumption of diffusion-limited forward rate constant being equal for all reactions only affects the results describing the kinetics of herd regulation. The concentrations of the complexes approach that of the equilibrium values regardless of the combinations of rate constants used. This is demonstrated in section 8.6 where we have studied the trajectories for the formation of *AB* and *AB_i_* when we fix the reverse rate constant to 10^-3^ s^-1^. These trajectories lack the initial transient peak for *AB_i_* but still converge to the same values as corresponding trajectories with a fixed forward rate constant.

In the last four sections, we have explored the effect of low-affinity interactors on tight binders and formulated a general model that describes herd regulation. In the next two sections, we have examined the effect of herd regulation in deterministic biological contexts and have explored its potential impact on binary outcomes.

### 1.2 Herd regulation in biological processes

Here we will illustrate how herd regulation could affect biological processes. In the first case, we will look at sex determination in *Drosophila melanogaster* where a two-fold increase in X chromosome-linked signal generates a binary response during the activation of the *Sxl* gene which determines the female identity. The second example is an illustration of a biological system of receptor and ligand that depend on threshold value determinants.

#### 1.2.1 Example 1: Low affinity binders are responsible for the absence of *Sxl* activation in *Drosophila melanogaster* males

Sexual identity in *Drosophila melanogaster* depends on the female-specific activation of the binary switch gene Sex-lethal (Sxl) before cellularization at cycle 14 (Cline, 1984). Sxl activation, in turn, is regulated by the Sxl establishment promoter *Sxl-Pe*. This process is dependent on X-linked signal elements (XSEs) and autosomal repressors. As females have two X chromosomes, they are able to produce twice the amount of XSEs as compared to males and can specifically activate *Sxl-Pe*. By contrast, *Sxl-Pe* remains off in males who only have one X chromosome. (In insects, the Y chromosome does not contribute to male determination unlike in mammals.). The molecular identities of XSEs are well known (Cline, 1988; Cline and Meyer, 1996; Erickson and Cline, 1993, 1991; Kappes et al., 2011; Keyes et al., 1992; Salz and Erickson, 2010). However, the precise mechanism underlying how the doubling of XSE induces the activation of *Sxl-Pe* remains unknown (Deshpande et al., 1995). Of note, no direct repressor that binds to *Sxl-Pe* during early nuclear cycles has been identified, thus far. Moreover, mutations in deadpan (*dpn*) and groucho (*gro*), two known negative regulators of *Sxl-Pe*, display relatively modest phenotypes (Chen and Courey, 2000; Lu et al., 2008; Mahadeveraju et al., 2020; Paroush et al., 1994; Younger-Shepherd et al., 1992).

Here we test the model that multiple low-affinity binders, capable of interfering with the interaction between activators and *Sxl-Pe*, function as regulators of *Sxl*. As the repression depends on several low affinity binders, it is relatively straightforward to explain why the forward genetic screens were unable to identify corresponding mutants. Single mutations typically isolated in the genetic screens will not display the requisite phenotype.

Our model posits that sex determination involves one tight binding complex, *AB*, regulated by weaker complexes *AB_i_* (refer to section 2.1.1, equations E1 and E2, and figure 4). Here, the DNA binding sites in *Sxl-Pe* and XSEs constitute tight binders (*A* and *B*) and form the activating complex (*AB*). We assume that binding of the low affinity repressors inhibits the promoter from binding to the activator, directly or indirectly. Low affinity interactors (*B_i_*) are repressors of *Sxl-Pe* that form the repression complex (*AB_i_*). We assumed that there are 50 binding sites in *Sxl-Pe* (Jinks et al., 2003), 200 XSE activators and 1000 low affinity repressors. The dissociation constant (*K_D_*) of the active complex is 10^-8^ M. The repression complex has a 100 fold lower dissociation constant (*K_D_i_*) of 10^-6^M. Here again, we considered identical forward reaction rates, whereas the reverse reaction rates are determined by the dissociation constant. Both XSEs and repressors compete for DNA binding sites in *Sxl-Pe*. We consider *Sxl-Pe* to be engaged in active transcription when 90% (45) of the binding sites are occupied by XSEs. Note that the values that we have used are estimations due to unavailability of biochemical characterization to obtain such values.

**Figure 4:**
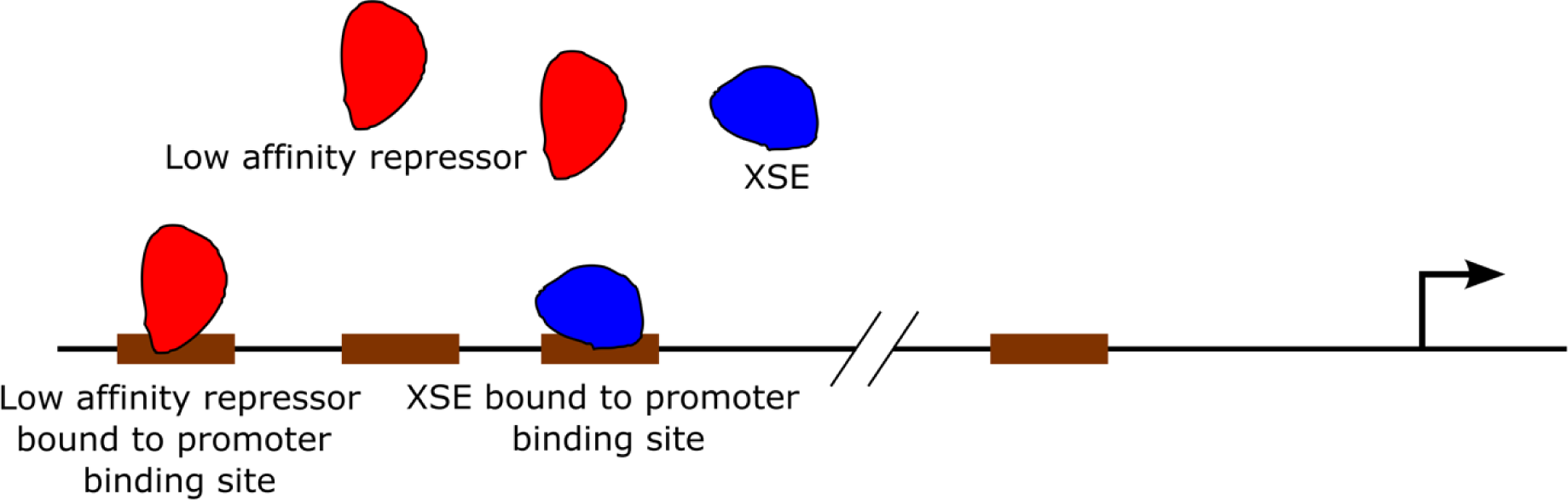
Components of our model of *Sxl-Pe* regulation. The blue objects represent XSEs while the red objects represent the low affinity repressors. The brown boxes represent protein binding sites on Sxl-Pe. The arrow depicts the transcription start site.

We used Gillespie stochastic simulations to study the kinetics of this system. We calculated the amount of XSEs and repressors bound to *Sxl-Pe* binding sites after 100 units of simulation time at different starting levels of XSEs and repressors. The simulation time scales are relative to the rate constants and may not correspond numerically to biological times.

We first assessed the activity of *Sxl-Pe* at different starting values of XSEs and repressors. As expected, the promoter is predominantly occupied by the repressors when activator levels are low. Increasing XSE levels elevates the number of activating complexes (figure 5A). Subsequently, we examined the effect of changing repressor levels on *Sxl-Pe* activity. Increasing the starting levels of repressors decreases the number of activator-bound *Sxl-Pe* sites. The binding sites are occupied by an equal number of activators and repressors when the starting levels of repressors are twice that of the starting levels of XSEs. This is despite a 100-fold difference in binding affinities (figure 5B).

**Figure 5:**
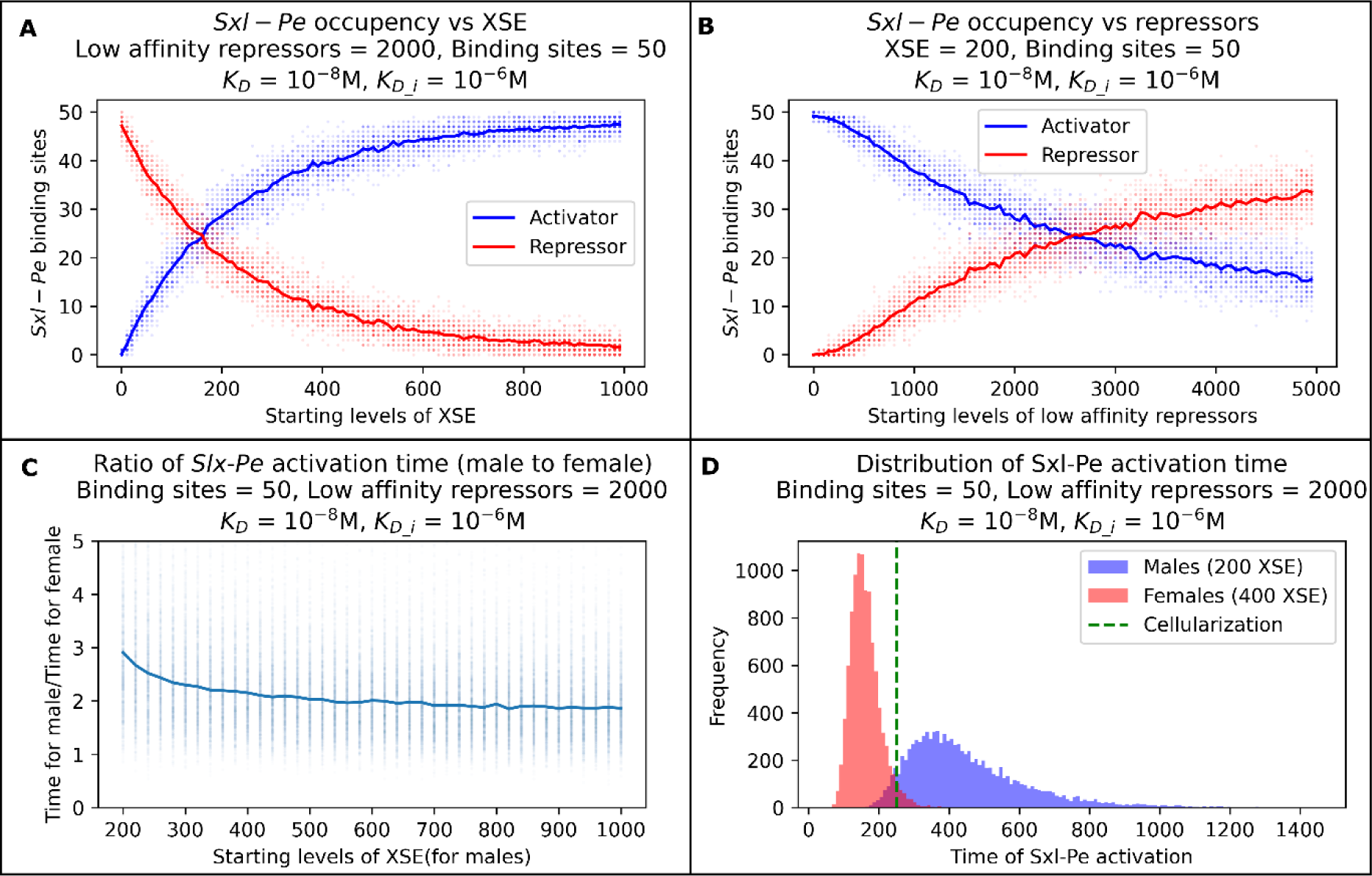
Model of sex determination with one set of low affinity binders. The amount of activating and repressing complexes formed when time = 100 units, at different starting values of XSEs and repressors (Panels **A** and **B** respectively). The ratio of time taken by males vs females to activate *Sxl-Pe* (XSEs occupy 90% of all available *Sxl-Pe* binding sites) (Panel **C**). Panel **D** shows the distribution of time taken for male and female cells to activate across a population of 10000 cells. The simulations were replicated 50 times for each data point in panels **A** and **B** (individual data points shown as scatter of the same colour), and 10000 for panel **C**.

Next, we focused on the time taken for the activation of *Sxl-Pe,* both in males and females. In wild-type *Drosophila melanogaster* embryos, activation of Sxl-Pe results in female fate whereas it remains off in males. Furthermore, this is a time-sensitive process as *Sxl-Pe* is active only between nuclear cycles 11-14 and the sex of the embryo is determined by early cellularization. (This is a narrow time window spanning just three nuclear cycles and takes about 30 minutes or ∼2000 seconds.). For this, we analyzed the ratio of time taken for *Sxl-Pe* activation in male nuclei and female nuclei in *Drosophila* embryos where sex determination occurs autonomously. The calculations accounted for the fact that the XSE levels in females are twice that in males. We found that this ratio is in a narrow range between 2 to 3 even when activator levels are varied between 200-1000 (figure 5C). Thus, activation of *Sxl-Pe* in males takes approximately twice as long as compared to females. We arbitrarily selected a test case where the starting level of XSE was 200 (for males) to study the time dependence of sex determination. We analyzed this over a population of 10,000 male and female simulations, each corresponding to a single nucleus. The time distributions in males and females for *Sxl-Pe* activation show two distinct peaks corresponding to the two sexes, with males (on average) taking appreciably more time for activation than females (figure 5D). Note that Sxl activation occurs over a range of time. Importantly, this pattern of nearly non-overlapping distributions indicates that within a given time interval, *Sxl-Pe* is activated only in females. There is however a small overlap between the male and female distributions (purple region in Figure 5D). This indicates that in a small subset of male nuclei, *Sxl-Pe* is turned on and vice versa in females. If the cellularization event occurs between the two peaks (green dotted line in Figure 5D), our model can temporally distinguish male and female *Sxl-Pe* activation. Thus, our simulations suggest that the time constraint imposed upon the activation of *Sxl*-*Pe* is significantly responsible for its female-specific activation. Furthermore, in the absence of such constraints, Sxl-Pe can be active even in male embryos, which has been demonstrated to lead to detrimental consequences (Cline and Meyer, 1996). Altogether, our analysis recapitulates both the sex-specificity and the temporal dynamics of *Sxl-Pe* activation. Increasing amounts of low affinity repressors increase the time taken for activation for both males and females. The effect of 1000 (half), 4000 (double) and zero low-affinity repressor on sex determination is shown in figure A2.

#### 1.2.1 Example 2: Signaling thresholds in biological systems can be explained by herd regulation

In general, molecular thresholds are crucial in several biological signalling events ranging from cell fate determination in early development to neuronal action potentials. Without such thresholds, signals resulting from stochastic noise (inherent to crowded cellular environments) would be able to aberrantly activate (or repress) biological processes. In this section, we describe a general molecular mechanism of how herd regulation can be instrumental in establishing such thresholds in biological systems.

Consider the example where receptor *A* is activated by binding a signalling molecule *B* to form the receptor-signal complex *AB*. The system also contains low affinity binders for the receptor and signalling molecule (*B_i_* and *A_i_* respectively) as we have explored in the previous sections. The receptor-signal complex is assumed to have a dissociation constant of 10^-8^ M whereas both the low affinity complexes have a dissociation constant of 10^-6^ M. We modelled the kinetics of this system using the Gillespie stochastic simulation with 50 receptors and 500 signalling molecules. This is in keeping with the observation that signalling molecules are usually in excess compared to the number of receptors (Cima, 1994). The system also contained 500 copies of each of the two low affinity binders. We monitored the levels of the receptor-signal complex (amount of *AB* formed) after 10000 units of simulation time, starting with different levels of the signal (*B*). The time of 10000 units was chosen to represent a phase of the reaction that is significantly distant from the fast-starting kinetics (see Figure A1A for a comparison of different time cutoffs). We simulated this system at different starting receptor levels and at levels of the low affinity receptors.

We observed a sigmoidal-like relationship between the number of receptor-signal complexes and the starting levels of the signal. When the number of signaling molecules in the system is below a threshold the rate of increase of the number of receptor-signal complexes is gradual. As soon as the number of signaling molecules exceeds the threshold the number of receptor-signal complexes increases sharply (sigmoid-like). Increasing the levels of the receptor molecule does not alter the threshold but only increases the final levels of the active receptor (figure 6A). Interestingly, we see that the threshold value is regulated by the number of low affinity receptors in the system (figure 6B). Whereas, varying the levels of low affinity signalling molecules does not affect the threshold (figure A1C). Nevertheless, the probability of collisions that lead to the tight binding complexes should approach zero on either increasing the low affinity signals or low affinity receptors. Without low affinity binders the relation between *B_0_* and *AB* is linear until *A* is exhausted (figure A1D). Thus, the signal threshold is dependent on the level of low affinity receptors in the system, whereas the amplitude of the signal is determined by the number of receptors.

**Figure 6:**
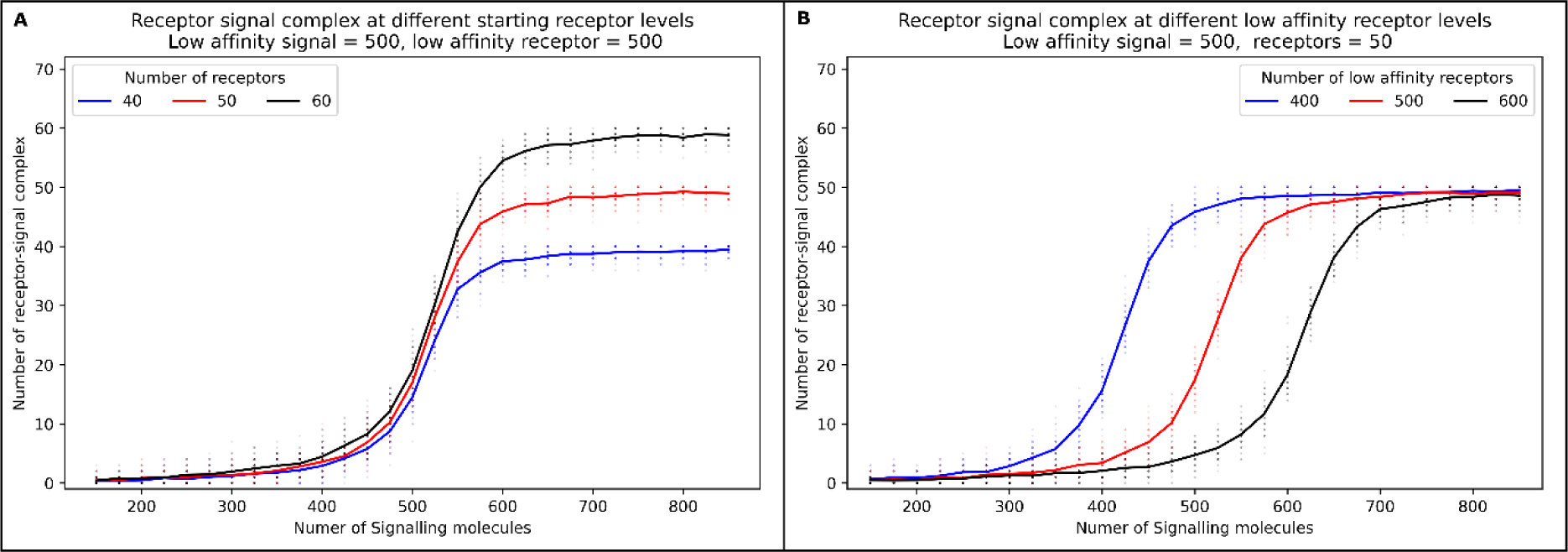
Modelling signalling thresholds in biological systems. All traces (averages of 50 replicates, individual data points shown as scatter) show the relationship between the active receptor and the number of signal molecules and low affinity complexes at different starting levels of the signal. Panel **A** shows this relation at three different starting receptor levels. Panel **B** shows this relation at different starting levels of the low affinity receptor. Plots are averages of 50 replicates and the scatter of the same colours shows the individual data points.

Iyer et al. (2022) analyzed the specification of different cell types under the influence of the Wingless morphogen gradient. They proposed that cellular compartmentalization and receptor promiscuity were critical to elicit a specific response to confer unique cell type identities. Interestingly, this was experimentally verified using the modulators of *Wingless* activity, *Dally* and *Dally like* (*B_i_*) which compete with *Frizzled-2* (*B*) receptor for *Wingless* (*A*) binding. The herd regulation model supports and extends these observations by providing straightforward mechanistic explanation for the observed phenomena. We also suggest the presence of undocumented low affinity binders of *Frizzled-2* (*A_i_*). In general, our model could be applied to a number of such biological systems where the outcome is contingent upon molecular thresholds.

In this section, we have investigated two biological scenarios where herd regulation might play a role. In the next section, we provide the design of an experiment which can validate the general concept of herd regulation.

### 1.3 Experiment to validate herd regulation based on DNA-DNA interaction

Our model of herd regulation holds for all types of molecular associations such as protein-protein, protein-DNA, DNA-DNA interactions etc. The binding affinity between single-stranded DNAs can be designed based on complementarity and melting temperature. Here we present the results of herd regulation in DNA-DNA interactions. In this example, each of *A*, *B*, *A_i_* and *B_i_* are single-stranded DNA (Figure 7). The sequences are such that the binding affinities are in the order, *AB* > *AB_i_*,∼= *A_i_B* > *A_i_B_i_*. Also, the sequences are designed to minimize self-complementarity and binding affinity of *AA_i_* and *BB_i_*.

**Figure 7:**
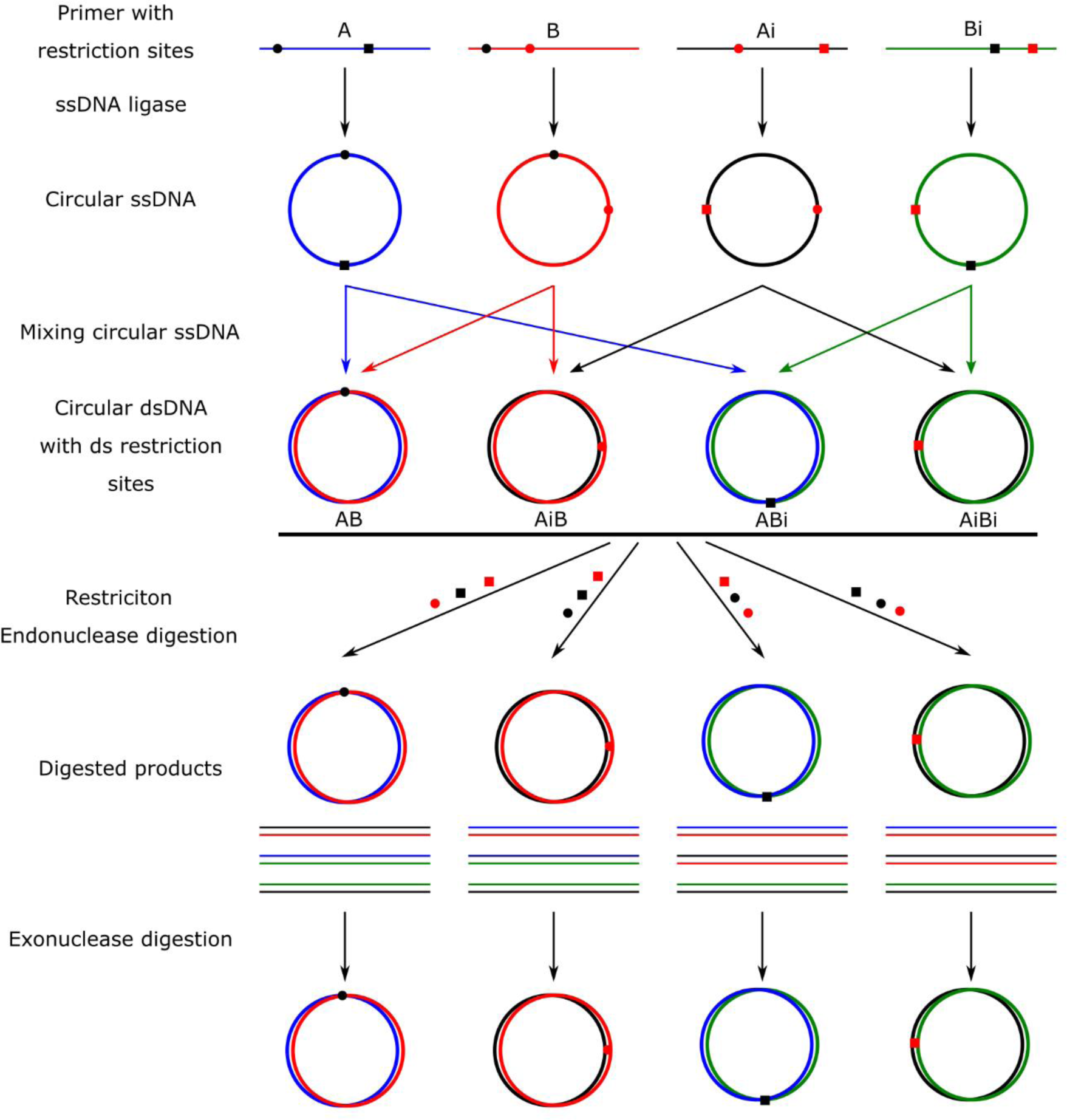
Schematic of the experiment to validate the effect of herd regulation. DNA strands are coloured blue, red, black, and green for *A*, *B*, *A_i_* and *B_i_*. red and black squares and circles represent different restriction sites and corresponding restriction enzymes. Different steps of the reaction are shown in the left-hand side.

The binding affinities between the species are controlled by mismatches between the sequences i) At the experimental temperature, the ΔG of binding of each combination corresponded to the binding energy. ii) Hybridization of ssDNA from two different species results in a unique restriction site (i.e. this location in the sequence has mismatches in all other combinations)

Let the unique restriction sites in *AB*, *A_i_B*, *AB_i_* and *A_i_B_i_* be, RE1, RE2, RE3 and RE4 respectively. Before hybridisation, each of the ssDNA (easily procured as oligonucleotides) is converted into circular ssDNA using a ssDNA ligase. These circular ssDNA which correspond to *A*, *B*, *A_i_* and *B_i_* are incubated together at the experimental temperature till equilibrium. After this, the reaction is quickly cooled to a temperature well below the lowest melting temperature to avoid any deviations from equilibrium. This should result in a mix of *AB*, *A_i_B*, *AB_i_* and *A_i_B_i_* products. To measure the concentrations of each product, four equivalent aliquots are made from this reaction mixture. To measure the amount of *AB*, the enzymes RE2, RE3 and RE4 are added to one of the aliquots to linearise double-stranded DNA corresponding to *A_i_B*, *AB_i_* and *A_i_B_i_*. The linear dsDNA is then removed by digestion using exonucleases. This is repeated for the other 3 products by using the enzyme combinations; RE1, RE3 and RE4 for *A_i_B*; RE1, RE2 and RE4 for *AB_i_* and RE1, RE2 and RE3 for *A_i_B_i_*. The products of this step contain the circular dsDNA of only one complex. DNA purification removes protein and buffer remains from enzyme reactions. The concentration of DNA is then measured using UV spectrophotometry at 260nm wavelength or by realtime PCR experiments.

This experimental method allows us to measure the steady state of the system after manipulating the concentrations and binding affinities of each of the reactants individually. Different reactions with different combinations of binding affinities and concentrations can be set up such that our results from steady-state (figures 1E, 2E and 6) can be replicated.

## 2 Discussion

Molecular investigations of biological processes almost always focus on interactions that are high affinity and specific. Seldom, if at all, is any importance given to interactions of low affinity/specificity. In part, this could be due to our inability to detect such interactions. In this study, we examine how the presence of low affinity interactors modulates the association of high affinity interactors. We term such modulation of tight binding events by low affinity interactors as ‘herd regulation’.

To illustrate herd regulation, we have considered a simple system of two interactors (*A* and *B*) that come together to form a dimeric complex (*AB*). The kinetics of AB formation was modulated in the background of low affinity interactors to both *A* and *B* (*B_i_* and *A_i_* respectively). In this case, we found that the rise time of the *AB* complex increases with changes in the levels of the low affinity interactors. Surprisingly, we find situations where *AB* is not the dominant product of the reaction, despite *K_D_* of *AB* several orders of magnitude higher. *A_i_* and *B_i_* when acting alone, without the presence of the other (section 2.1.1), are weak binders and offer weak competition. Only when acting in consort (section 2.1,2) do they become potent competitors. This can be explained by the lack of monomeric *A* and *B* in the system, as all available *A* and *B* are consumed by the formation of *AB*, *A_i_B* and *AB_i_*. This is the case since *A* and *B* contribute to the formation of two complexes each, whereas *A_i_* and *B_i_* are a part of only one complex each. Thus, the products of the dissociation of *A_i_B* or *AB_i_* have a higher probability of encountering *A_i_* or *B_i_* compared to *A* or *B*. This study effectively predicts the ranges of both the concentrations and dissociation constants at which herd regulation would be effective. In general, herd regulation is affected by the quantity of the low affinity binders as well as their respective binding efficacies (quality). Interestingly, our data argues that the quality is more impactful than the quantity. We successfully reproduce all of these results almost exactly using a separate continuous method.

In our simulations, we have demonstrated herd regulation of a binary interaction of *A* and *B* (giving *AB*) with interference from low affinity competitors *A_i_* and *B_i_*. In the crowded cellular environment, reactions could be more complex, involving multiple reactants with varying stoichiometry. It is thus relatively straightforward to imagine that herd regulation would impact such complex interactions more than it does the simple *AB* system. The other important aspect here is that the low affinity competitors could be multiple species. For instance, the interactor *A* could have many homologous or structurally analogous molecules in the cell, all of which could collectively constitute the *A_i_*. Given the sizes and abundances of protein families, and the comparatively low number of folds, our estimates of the levels of the low affinity competitors are conservative. Please note that similar arguments would also hold for herd regulation involving different types of molecules such as DNA, RNA etc.

Unlike our simulations, in a cell all the potential interactors may not coexist or be active at the same time. It is conceivable that before one of the strong binding interactors is introduced into the system, the other binding partner and the low affinity interactors are already present. Such a situation would only magnify the effects we have discussed above. i.e., the relative rates of reactions could be slower than what we have simulated, and the time taken to equilibrate the system would be longer. For instance, during sex determination in *Drosophila*, the DNA binding sites in *Sxl-Pe* exist in the cell even when the XSEs are absent. In this situation, it is reasonable to assume that these DNA binding sites are bound to a cohort of low affinity repressors. The XSEs produced during early development (nuclear division cycles 7-13) will have to displace the low affinity binders for the DNA binding site to activate *Sxl*.

In our computations that study kinetics, we have assumed that the forward rate constants (*k_forward_* or *k_f_*) for the high and low affinity interactors are the same. This is not an unusual circumstance in biological processes. For instance, proteins that recognize specific sequences of DNA initially bind non-specifically and diffuse along the DNA to its cognate binding site (Kim and Larson, 2007; Shimamoto, 1999; Wang et al., 2006). Even, non-specifically binding proteins would also have a similar rate of association with the DNA as specific binders (Cima, 1994; Schreiber et al., 2009). The rates of dissociation, however, are vastly different and determine the binding affinities. A similar argument can be made regarding protein-protein (or protein-peptide) and protein-RNA interactions, especially if the binding was predominantly due to complementarity in electrostatics. In the studies of equilibrium/steady state this assumption of comparable forward rate constants is not required. This is because the equilibrium state is only dependent on the dissociation constant and starting concentrations of the reactants.

It should be noted that the herd regulation model presented in this study is basic. We built such a model to set up a platform where one could build in additional constraints for future investigations. Such future directions include: i) The effect of herd regulation on oligomerization and/or polymerization reactions; ii) Our models demonstrate regulation when there is a fixed quantity of the reactants. A more realistic model would also incorporate rates of their production and degradation; iii) Modeling interactions where there are transition states and more involved kinetics. iv) Proving herd regulation experimentally would constitute a significant challenge as low affinity interactors are mostly undetected. We have addressed this by suggesting an *in vitro* experiment involving circular single stranded DNA. We believe that these experiments could validate the findings reported in results section 2.1. We also have a suggestion for *in vivo* investigation (see below). The present study lays down the basis to investigate such questions, which is otherwise beyond the scope of this study.

We have illustrated herd regulation in biological context with two examples and also suggest one ‘in vitro’ experimental stratergy to verify the model. In one example, we use protein-protein interactions akin to signal-receptor activation and in the other, we use DNA-protein interactions to illustrate sex determination in *Drosophila melanogaster*.

In the example of signalling, we resort to our simple *AB* model (*A* = receptor, *B* = signal or a ligand). The signal (*B*) can bind its cognate receptor *A* or any of the low affinity competitive/homologous receptors *A_i_*. Once *B* is at a level (threshold) where it has saturated all the *A* and *A_i_*’s the kinetics is only dependent on the level of *B_i_* (low affinity competing signals). Thus, exhaustion of *A_i_* switches the system, from herd regulation influencing both the tight binders, to one where it affects only a single tight binder. Thus, the threshold is primarily dependent on the amount of *A_i_*, the low affinity receptor, present in the system. This formulation also helps explain earlier observations and computations where the presence of promiscuous receptors (*A_i_*) determines cell fate in *Drosophila melanogaster* development (Iyer et al., 2023). Note that our model is significantly simpler than the model proposed by Iyer et al which required invoking compartments and significantly more rate equations to explain this phenomenon. We can explain this with three simple equations (E9, E10 and E11) (section 2.1.4 and appendix section 8.4). This also simplifies the evolution and design of molecular switches. Consider *A*, *B*, *A_i_* and *B_i_* where *A* and *B* interact strongly whereas *A_i_* and *B_i_* are homologs of *A* and *B* which have partially similar binding sites and thus interact with *A* and *B* and with each other. This system will become a molecular switch if there are mutations (engineered or evolved) in the binding interface such that *A_i_B_i_* is not formed (see sections 2.1.2 and 2.1.3).

In the example of sex determination, we analysed a binary switch gene, *Sxl*, whose activity confers female identity. *Sxl* activity is primarily affected by the presence of X-linked signalling elements (XSEs) and autosomal repressors. XSEs are derived from the X chromosome. Thus, females with two X chromosomes activate *Sxl-Pe* while the *Sxl-Pe* of males remains inactive. Sex determination is dependent on the activity of *Sxl-Pe* at the end of the syncytial blastoderm stage of embryogenesis (NC10-13). We have modelled sex determination considering the binding sites in *Sxl-Pe* as *A*, XSEs as *B* and the autosomal repressors as low affinity interactors, *B_i_*. Note that many of the *Sxl-Pe* binding sites are concentrated within 400 base pairs in the promoter. (Estes et al., 1995). We found that the presence of low affinity binders temporally resolves *Sxl-Pe* activation between males and females. XX and XY genotypes would determine female and male states respectively as long as the cellularization occurs at a time point between male and female activation. The primary factor that allows for this temporal decision is the amount of XSEs, the cell cycle at which cellularization happens and the amount of low affinity autosomal repressors present in the system. Among these, the amount of XSEs and timing of cellularization has been experimentally perturbed. Mutations in XSE genes are detrimental to females (Cline and Meyer, 1996). Delayed cellularization generates embryos with more *Sxl-Pe* activation and vice versa (Erickson and Quintero, 2007). It would be interesting to see the effect of perturbing the low affinity binder levels *in vivo*, although this would require the identification of multiple low affinity binders which would need to be mutated simultaneously. Hypothetically, changing the level of the low affinity repressors (herd regulators) could affect the male-female ratio in a population (see figure A3 for distributions of *Sxl-Pe* activation at different low affinity repressor levels). Since the sex determination decision is made individually in each nucleus in an autonomous manner, this effect should manifest as a mosaic phenotype as those shown in (Erickson and Quintero, 2007). The presence of such repressors is our prediction. The role of deadpan (*dpn*) and groucho (*gro*) as weak negative regulators is already reported (Chen and Courey, 2000; Lu et al., 2008; Mahadeveraju et al., 2020; Paroush et al., 1994; Younger-Shepherd et al., 1992). However, *gro* does not bind DNA directly and needs to be recruited to the promoter and neither deadpan nor *gro* active in all the cells (sex determination decision happens in all the cells). We speculate that the unknown entities responsible for the recruitment of *gro* along with other yet unknown entities have a combined ‘herd’ repressive effect on the promoter. Our model of sex determination can be validated only after these low affinity repressors are identified.

It will be important to consider these data in a broader, more general context. Even a cursory examination should reveal that binary decisions constitute important cornerstones of biological systems both at the ‘micro’ and ‘macro’ levels. It is noteworthy that many of the early foundational discoveries, especially in the field of developmental genetics were aimed at uncovering the presence of a single master determinant that functions as a molecular switch. (Sxl is just one such example of the several well documented ‘master switch genes’). As the proper execution of the downstream molecular processes is critically dependent on the early choice, these binary switches have proven to be highly effective means to regulate major outcomes of a variety of determinative events. Notwithstanding the benefits however, one ought to consider a few important caveats that are readily overlooked while considering biological relevance and regulation of binary mechanisms that underlie fate determination. The genetic and biochemical strategies aimed at the identification and characterization of canonical bio-switches (and their regulatory components) invariably rely on phenotypes that are extreme. For instance, the genetic screens that identified regulators of sex determination relied on either male or female specific lethality. (The nomenclature of the relevant mutations *daughterless*, *sisterless a, b* etc. effectively underscores this feature). As the detection of relevant mutations relied on highly penetrant sex-specific lethality, a number of potentially interesting albeit weak mutations were likely dismissed. Their detection is only possible upon simultaneous deletion or addition of multiple weak affinity factors as approximated in our computational approach.

We have also observed an overlap in the distribution of time taken for male and female *Sxl* activation. This suggests that binary switch systems are not strictly binary. Their outcome is actually two states that in many cases can have small (but not insignificant), overlaps. This becomes especially important in the case of cell autonomous determinative events. For instance, the sex of every single cell in a *Drosophila* embryo is independently determined by the total concentration of X-chromosomal elements. As a result, gynandromorphs consist of cells of both sexual identity and, on rare occasions can survive even up to adulthood. While every instance does not result in a dramatic and demonstrable outcome such as this, it is conceivable that several cells of the ‘wrong sex’ may be determined in an organism and are eventually eliminated (or rendered ineffective) without any detectable consequence. Analogous scenarios can be envisaged in other biological contexts i.e., a few haltere-like cells could co-exist in wings or some neurons may exhibit partial epithelial traits. Such heterogeneity has been well documented in primordial germ cells in blastoderm stage *Drosophila* embryos while its relevance remains to be determined.

Canonical models suggest that both the initiation and maintenance of the binary switch genes follow a stereotypical path toward a unified outcome. Altogether, while binary decisions may lead to invariable and robust outcomes at a macro-or organismal level, cellular heterogeneity likely exists at a micro-or cellular level. Mechanisms ought to be in place that do not allow such variability to exceed permissible limits leading to detrimental biological consequences. Importantly, the tolerable threshold may differ vastly depending on the cellular or organismal contexts. The mechanistic underpinnings and biological relevance of such mechanisms need closer scrutiny and careful consideration as these represent molecular hurdles that are likely essential components of homeostatic mechanisms.

## 4 Materials and Methods

### 4.1 Reactions

Consider the following reaction,

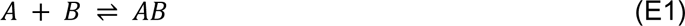

Consider the following reactions happening in proximity to E1,

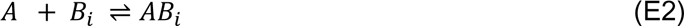

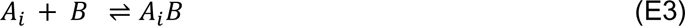

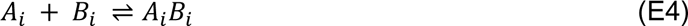

Here *A*, *B*, *A_i_* and *B_i_*, each represent ensembles of biomolecules. The Interactions between any pair of *A* and *B* have high affinity whereas the interaction between *A_i_* and *B_i_*, have low affinity. Note that although these interactions are low affinity, they are specific.

### 4.2 Gillespie stochastic simulations

Differential equations can describe the progression of reactions deterministically. But they assume that the reaction volume is infinitely large, and the stochastic behaviour of molecules is cancelled out. This also means that the concentration terms are continuous. But a cell is a small system with relatively few molecules. In such a system the concentration is no longer continuous, and randomness could often play an important role. We use the direct method of the Gillespie stochastic simulation algorithm (Gillespie, 2013, 1977) to simulate such a system. This is a numerical method for simulating the temporal evolution of systems of chemical reactions with randomness. It involves iteratively sampling reaction events and updating the state of the system based on the chosen reaction and the current state of the system. And thus can be used to study the behaviour of complex chemical systems over time, including the effects of noise and uncertainty on the system’s dynamics.

We implemented the Gillespie algorithm, in python3. First, the initial state of the system is set, including the number and types of reactants and the reaction rates for each chemical reaction. Then, for each possible reaction, the propensity, a measure of the likelihood that the reaction will occur in the next time step, is calculated based on the reaction rate and the current state of the system. *A* random number generator is used to choose one of the possible reaction events based on the propensities calculated. The state of the system is then updated by adding or subtracting reactants as appropriate. The simulation time is advanced by a time step calculated based on the inverse of the sum of propensity and a random number. The process is repeated until the system reaches a) a pre-specified time b) a specified number of iterations c) when a product crosses a pre-specified level, or d) steady state (described in section 4.2.2).

#### 4.3.1 Parameters used to generate the results

Section 2.1.1: *A_0_* = 500, *B_0_* = 500, *K_D_* = 10^-8^ M and *C_0_* = 10^-5^M and *N_0_ =* 1000. *B_i0_* and *K_D_i_* were varied in the section.

Section 2.1.2: *A_0_* = 500, *B_0_* = 500, *K_D_* = 10^-8^ M and *C_0_* = 10^-5^M and *N_0_ =* 1000. *A_i0_*, *B_i0_* and *K_D_i_* were varied in the section. Note that in all the cases *A_i0_* = *B_i0_* and their binding affinities were also kept the same and represented by the common term *K_D_i_*.

Section 2.1.3: *A_0_* = 500, *B_0_* = 500, *A_i0_* = 5000, *B_i0_* = 5000, *K_D_* = 10^-8^ M, *K_D_i_* = 10^-6^ M and *C_0_* = 10^-5^M and *N_0_ =* 1000. *K_D_cross_* was varied in the analysis.

Section 2.2.1: *A_0_* = 50, *B_0_* = 200, *B_i0_* = 2000 and *K_D_* = 10^-8^ M, *K_D_i_* = 10^-6^ M and *C_0_* = 10^-5^M and *N_0_ =* 1000. *B_i0_* and *B_i0_* were varied in the section.

Section 2.2.2: *A_0_* = 50, and *K_D_* = 10^-8^ M, *K_D_i_* = 10^-5^ M and *C_0_* = 10^-3^M and *N_0_ =* 100.. A*_0_*, *B_0_* and *B_i0_* were varied in the section. The simulations were carried out till the simulation time of 10000 units.

Also, see section 4.3.3 and appendix 8.3 for the justification for the values provided.

#### 4.3.2 Steady state calculations

For reactions with only *B_i_* present, steady-state was assumed to have been reached when the average fluctuation over 1000 steps for every reactant and product is less than 1 (molecule or molecular complex). This was evaluated in intervals of 100 steps.

For reactions with both *B_i_* and *A_i_,* steady-state was assumed to have been reached when the average fluctuation of over 1,000,000 steps for every reactant and product is less than 1 (molecule or molecular complex). This was evaluated in intervals of 100,000 steps.

#### 4.3.3 Calculation of propensities

The propensities were calculated using the following equations

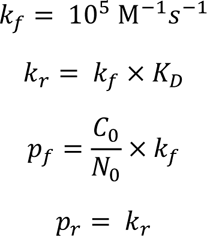

Where *k_f_* and *k_r_* are the forward and reverse rate constant respectively. *k_f_* is diffusion controlled and set at 10^5^ M^-1^s^-1^ (Schlosshauer and Baker, 2002).*p_f_* and *p_r_* are the forward and reverse propensities used in the Gillespie stochastic simulations. *C_0_* and *N_0_* are used to scale the second-order forward rate constant (*k_f_*) based on concentration.

For sections 2.1 and 2.2.1, *C_0_* = *10^-5^M* and *N_0_ = 1000.* For section 2.2.2 *C_0_ = 10^-3^M* and *N_0_ = 100* (Traces for other values of *C_0_* can be found in figure A1B) (justifications for the values are provided in appendix section 8.3).

### 4.4 Parameters for generating results from differential equations

The parameters used are the same as those described in 4.3.1, except that the number of molecules were converted into concertation using the following formula based on effective concentrations of the number of particles in the Gillespie simulations, calculated as follows.

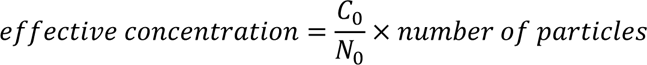

Initial concentration of Ai is set to 0 for calculations in sections 8.4.1. The K_Dcross_ for the dissociation of AiBi is set to infinity in all sections except section 8.4.3.

Trajectories were calculated by Runge-Kutta 4th Order numerical integration. The equilibrium states were calculated by solving equations E10, E11 and E12 for *AB*, *AB_i_* and *A_i_B*. The system of equations was solved by sympy module in python3. Real solutions that satisfy stoichiometry are selected.

### 4.5 Data availability

All scripts/programs used in this study along with the data they have produced can be accessed at http://sites.iiserpune.ac.in/~madhusudhan/Herd_regulation/figures.tar.gz

## Acknowledgements

We would like to thank - Dr. Neelesh Soni and Aditya Marodia for help with the Gillespie algorithm and debugging scripts, Dr. Chetan Gadgil and Dr. Anirban Hazra for critical reading and criticisms of the manuscript and COSPI lab members. The support and the computational resources provided by the PARAM Brahma facility under the National Supercomputing Mission, Government of India at the Indian Institute of Science Education and Research (IISER) Pune are gratefully acknowledged. SM would like to acknowledge CSIR SRF fellowship. This work was supported by the Department of Biotechnology, Government of India grant to the Indian Institute of Science Education and Research Pune; Department of Biotechnology, India under BICB center grant (BT/PR40262/BTIS/137/38/2022).

## 8 Appendix

### 8.1 Effect of changing levels of low affinity signaling molecules on the threshold

**Figure A1:**
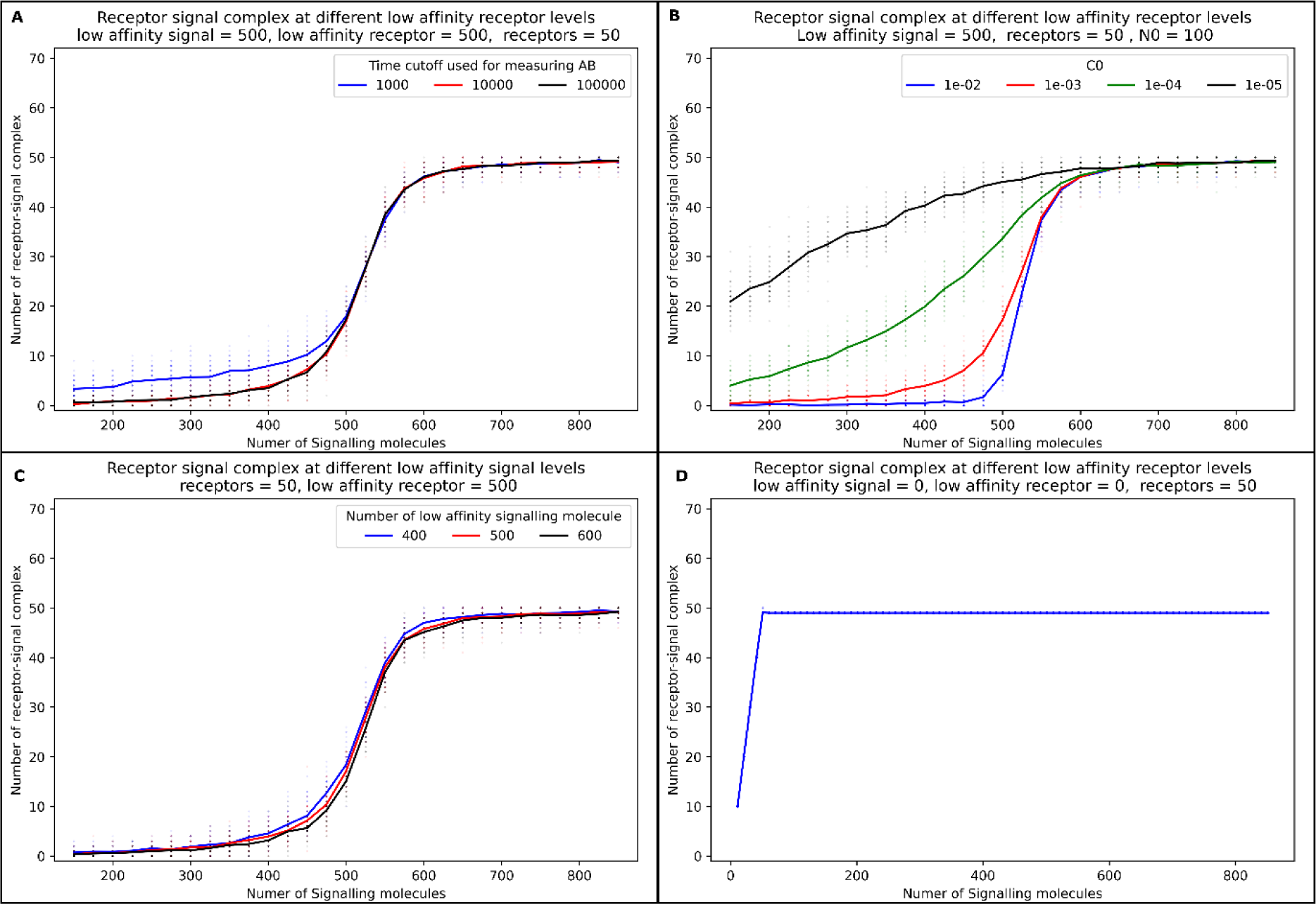
Dependence of receptor activation in varying levels of decoy signalling molecules in panel A, different *C_0_* values in panel B and different time cutoffs used for sampling in panel C. Panel D shows a control case of herd regulation when there is no low-affinity binder present. The difference in the scale of x-axis was required show show the the results between 0-50. In the range of A, B and C this plot would be a flat line with saturated AB. The parameters used were the same as that in Figure 5.

### 8.2 Male and female Sxl-Pe activation at different starting level of low affinity repressors

**Figure A2:**
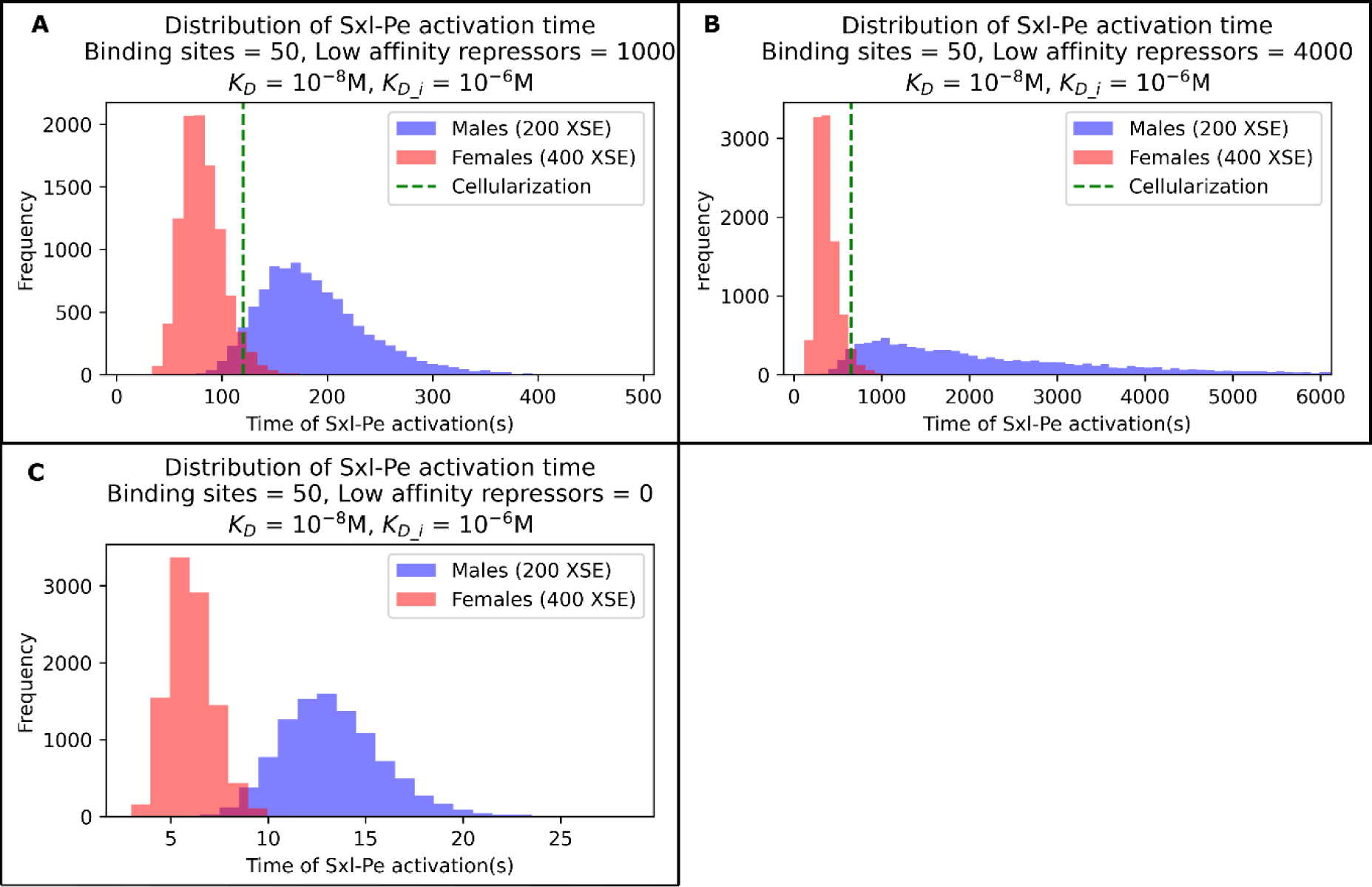
Effect of low affinity repressors on male and female activation times. Panel A shows male and female activation when the starting levels of low affinity repressors are 1000 (half that of figure 5D). Panel B shows the effect of 4000 (double that of figure 5D) low affinity repressors on male and female activation. Please not the variation in scales of x axis.Panel C shows male and female activation when there are no low affinity binders present. Parameters are shown in the top of the plots.

### 8.3 Local concentrations of reactants

Here we are addressing the physiological relevance of the concentrations of interactors used in the study. To do this, we estimate the volume that contains a single interactor. This in turn gives us the distance that separates two interactors.

The average volume (m^3^) occupied by one interactor is calculated from the concentration as

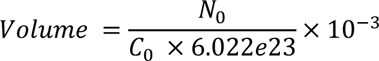

Where *N_0_* is the number of interactors and *C_0_* is the concentration. 6.022e23 is Avogadro’s number and 10^-3^ is the conversion factor from dm^3^ to m^3^.

Assuming that the volume containing the interactor is spherical

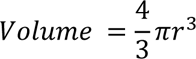

Where *r* is the radius of the sphere. The average distance (*d*) that separates two interactors is 2*r*

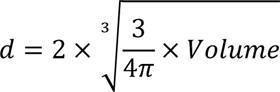

In this study, we have considered reactant concentrations ranging from 10^-5^M to 10^-3^M. We have also worked with the number of interactors in the range from (of the order of) 100 – 1000. Substituting these values in the equation above gives us interactor distance ranging from 68.2nm to 682 nm. Considering that the sizes of these interactors are of the order of a few nm and also that cellular and sub-cellular compartments are micrometre scale objects (Maul and Deaven, 1977), the concentrations do not represent overly crowded or unphysical scenarios. In fact, in several instances, the concentrations of cellular interactors are shown to be much higher than our somewhat conservative estimates.

Regulatory processes in biological systems usually occur under conditions of local enrichment, e.g. localization in subcellular compartments (Roth et al., 1989), nuclear speckles (Chen and Belmont, 2019) and lipid rafts in the cell membrane (Simons and Ehehalt, 2002). Such local enrichments have been shown to increase effective concentrations by as much as 5 orders of magnitude (Abrami and van Der Goot, 1999). In comparison, the values of concentrations used in this study are conservative.

### 8.4 Kinetics and steady state of herd regulation can be replicated by differential equations

#### 8.4.1 *B_i_* affect the formation of *AB*

**Figure A3:**
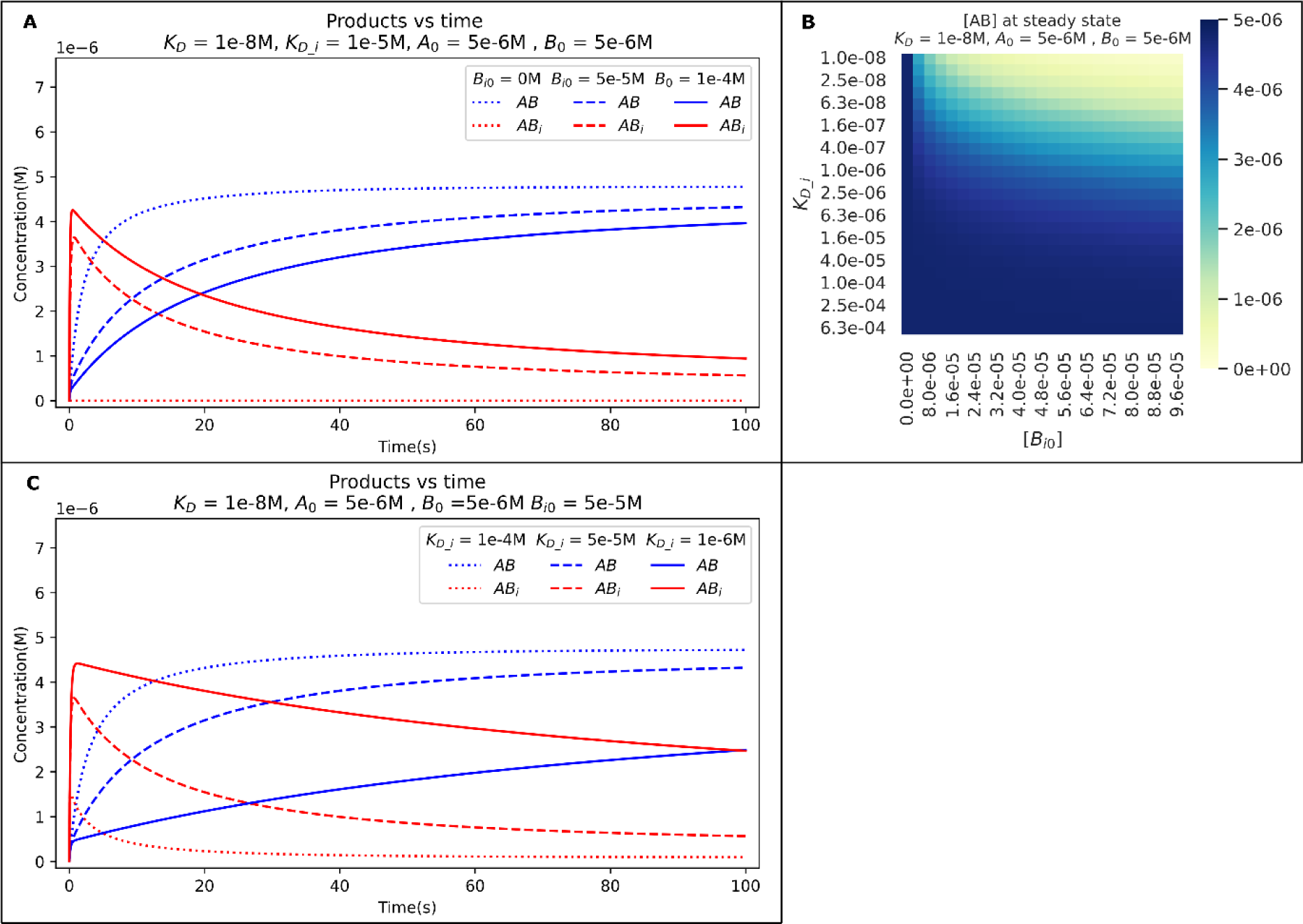
Panel A shows the effect of different concentrations of *B_i_0* on *AB* formation. Panel B shows the concentrations of *AB* at different levels of *K_D_i_* and *B_i_0*. Panel C shows the effect of *K_D_i_* on *AB* formation. The concentration scales is noted in the top left corner of the plot

#### 8.4.2 *B_i_* and *A_i_* affect the formation of *AB*

**Figure A4:**
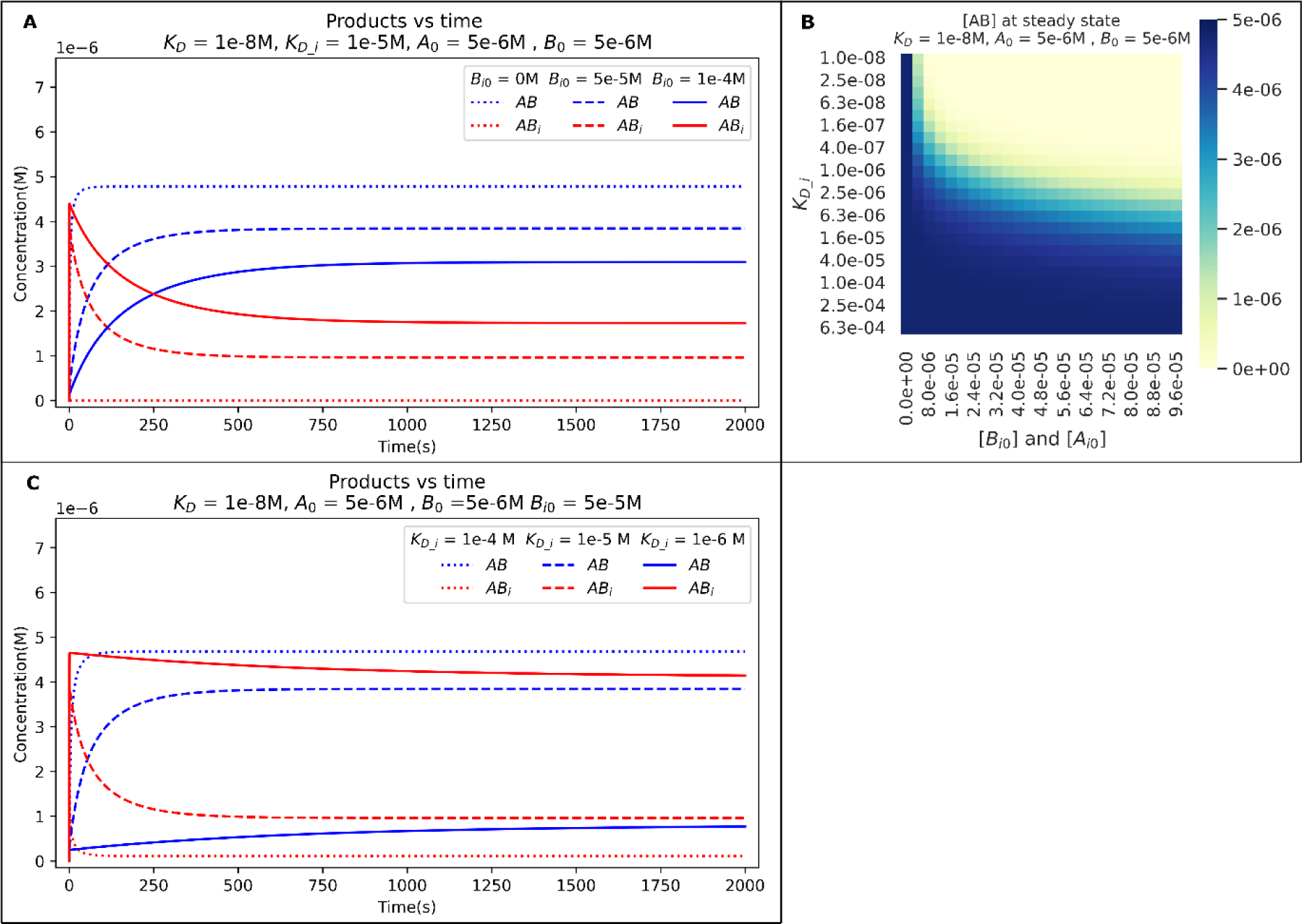
Panel A shows the effect of different concentrations of *B_i_0* on *AB* formation. Panel B shows the concentrations of *AB* at different levels of *K_D_i_* and *B_i_0*. Panel C shows the effect of *K_D_i_* on *AB* formation. Parameters used are shown above the graph. The concentration scales is noted in the top left corner of the plot

#### 8.4.3 Effect of cross reaction between *B_i_* and *A_i_* on the formation of *AB*

**Figure A5:**
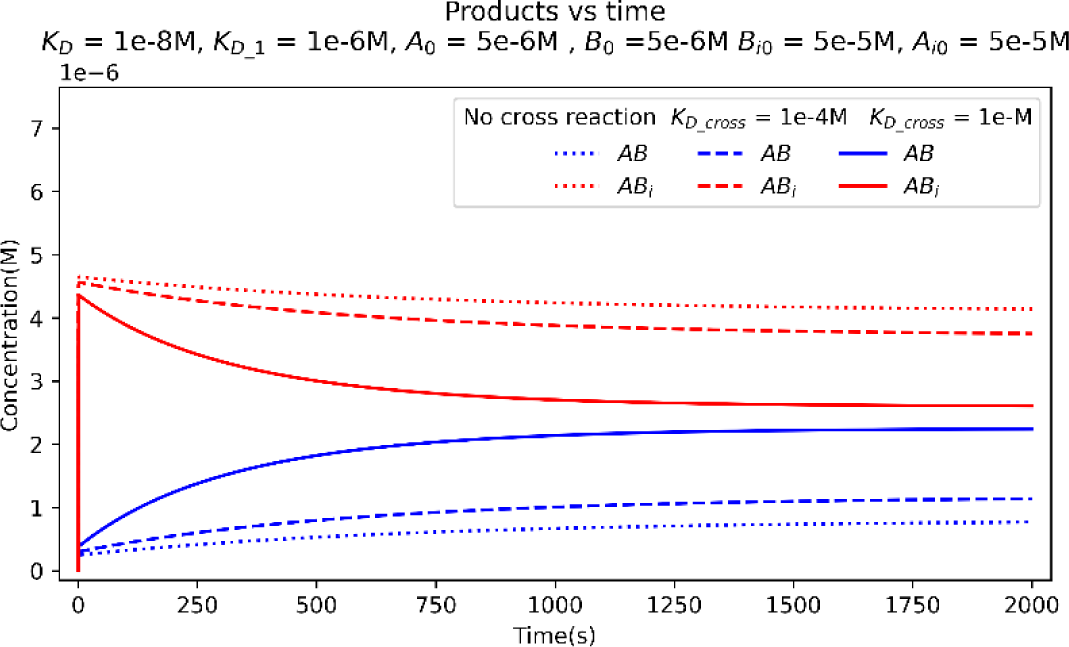
Figure shows the effect of cross reaction on AB formation.

### 8.6. Effect of herd regulation in higher order when reverser rate constants are fixed

**Figure A7:**
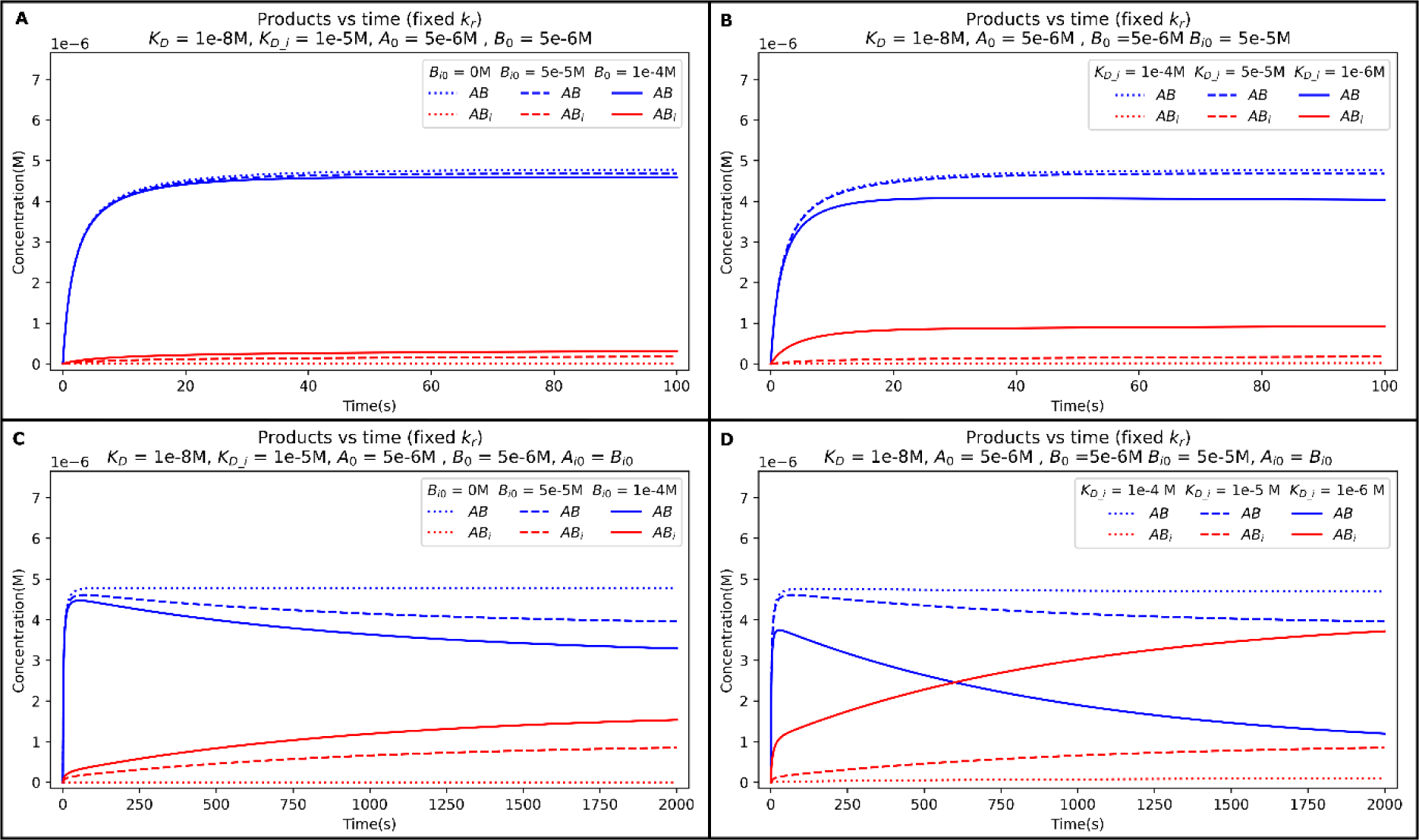
Trajectories with fixed reverse rate constant. The reverser rate constant for all the reactions in this plot are set to 10^-3^ s^-1^. The equivalent trajectories of reactions show in figure A3 A and C are shown in panels A and B and A4 A and C are shown in panels C and D. The parameters used to plot the graphs are mentioned in the in the respective graphs.

